# Invasive *Aedes (Fredwardsius) vittatus* reaches Continental America

**DOI:** 10.1101/2025.10.29.684036

**Authors:** Rahuel J. Chan-Chable, César R. Rodríguez-Luna, Román Espinal-Palomino, Carlos N. Ibarra-Cerdeña

**Author notes:** Address for correspondence: Carlos N. Ibarra-Cerdeña, Human Ecology Department, Center for Research and Advanced Studies (Cinvestav), Carretera Mérida-Progreso, Loma Bonita, 97205 Mérida, Yucatán, México.

## Abstract

We report the first population of *Aedes* (*Fredwardsius*) *vittatus* in continental America, detected in Yucatán, Mexico. Phylogenetic analysis clustered the Mexican sequence with Caribbean lineages (posterior probability 0.8–0.9), suggesting introduction via the Caribbean. Given its arbovirus competence, urgent inclusion in surveillance programs is warranted.

---

Mosquito-borne arboviruses such as dengue, Zika, chikungunya, and yellow fever have expanded dramatically over the past five decades, driven by urbanization, globalization, and human mobility (1). Dengue and chikungunya alone now cause more than 50 million infections annually, reflecting a thirty-fold increase linked to demographic and ecological change (2). While *Aedes aegypti* and *Aedes albopictus* remain the primary invasive vectors under surveillance and control, other species of epidemiological relevance are gaining increased attention as potential emerging threats (3).

*Aedes* (Fredwardsius) *vittatus* (Bigot, 1861) is one such mosquito, notable for its expanding range and proven arboviral vector competence (*4*). Described from Corsica, France (*5*), *Ae. vittatus* is now distributed across Africa, the Mediterranean Basin, the Middle East, and South and Southeast Asia, with sporadic detections in southern Europe and the Caribbean. The species is highly adaptable, breeding in both natural and artificial containers, and thrives in sylvatic, rural, agricultural, and peri-urban environments (6). Laboratory and field studies confirm its ability to transmit dengue, chikungunya, Zika, and yellow fever viruses, with additional potential for Japanese encephalitis and West Nile (7). Its recent detection in continental America, specifically in Mexico’s Yucatán Peninsula, highlights both its ecological plasticity and the urgent need to investigate introduction pathways and its potential role in arboviral transmission.

During entomological surveillance in August–September 2025, we collected 67 adult *Aedes* (Fredwardsius) *vittatus* in traditional Mayan cornfield (milpa) (Appendix figure 1) on the outskirts in the municipalities of Mama and Teabo, Yucatán, Mexico (Table 1; Figure 1A). Adults were aspirated as they attempted to bite field personnel (Figure 1B; Table 1) Both sexes were present (Figure 1C-D), supporting evidence of local reproduction and establishment in rural agricultural environments. Specimens were morphologically identified using standard taxonomic keys (4,5), and vouchers were deposited in the Arthropod Collection (ECO-CH-AR), ECOSUR, Chetumal Unit. *Ae. vittatus* can be distinguished from other *Aedes* species by its dark proboscis with pale yellowish scales, small bilateral patches of white scales on the clypeus, three pairs of narrow white patches on the anterior scutum, a short maxillary palp with apical white scaling, and a distinct white patch at the midpoint of the third tibia (Figure 1C-F).

**Table 1.** Collection records of *Aedes* (*Fredwardsius*) *vittatus* in the Yucatan Peninsula, southeastern Mexico, documented in this study.

| Collection date | Collection time | Location | Municipality | State | Latitude (N) | Longitude (W) | Elevation (m.a.s.l.)<sup>1</sup> | No. Mosquitoes |
| --- | --- | --- | --- | --- | --- | --- | --- | --- |
| 24/08/2025 | 16:00 h | Mama | Mama | Yucatán | 20.480198° | -89.357942° | 23 | 2 Females |
| 24/08/2025 | 16:30 h | Mama | Mama | Yucatán | 20.480300° | -89.357940° | 22 | 1 Male |
| 25/08/2025 | 06:00-07:00 h | Mama | Mama | Yucatán | 20.480049° | -89.357824° | 23 | 4 Females |
| 06/09/2025 | 16:00-18:00 h | Mama | Mama | Yucatán | 20.480198° | -89.357942° | 23 | 3 Females |
| 06/09/2025 | 16:00-18:00 h | Mama | Mama | Yucatán | 20.480198° | -89.357942° | 23 | 5 Males |
| 17/09/2025 | 16:00-18:00 | Mama | Mama | Yucatán | 20.480049° | -89.357824° | 23 | 33 Males |
| 17/09/2025 | 16:00-18:00 | Mama | Mama | Yucatán | 20.480049° | -89.357824° | 23 | 18 Females |
| 17/09/2025 | 16:00-16:30 | Teabo | Teabo | Yucatán | 20.40315° | -89.287062° | 22 | 1 Male |

<sup>1</sup> m.a.s.l., meters above sea level.

**Figure 1.**
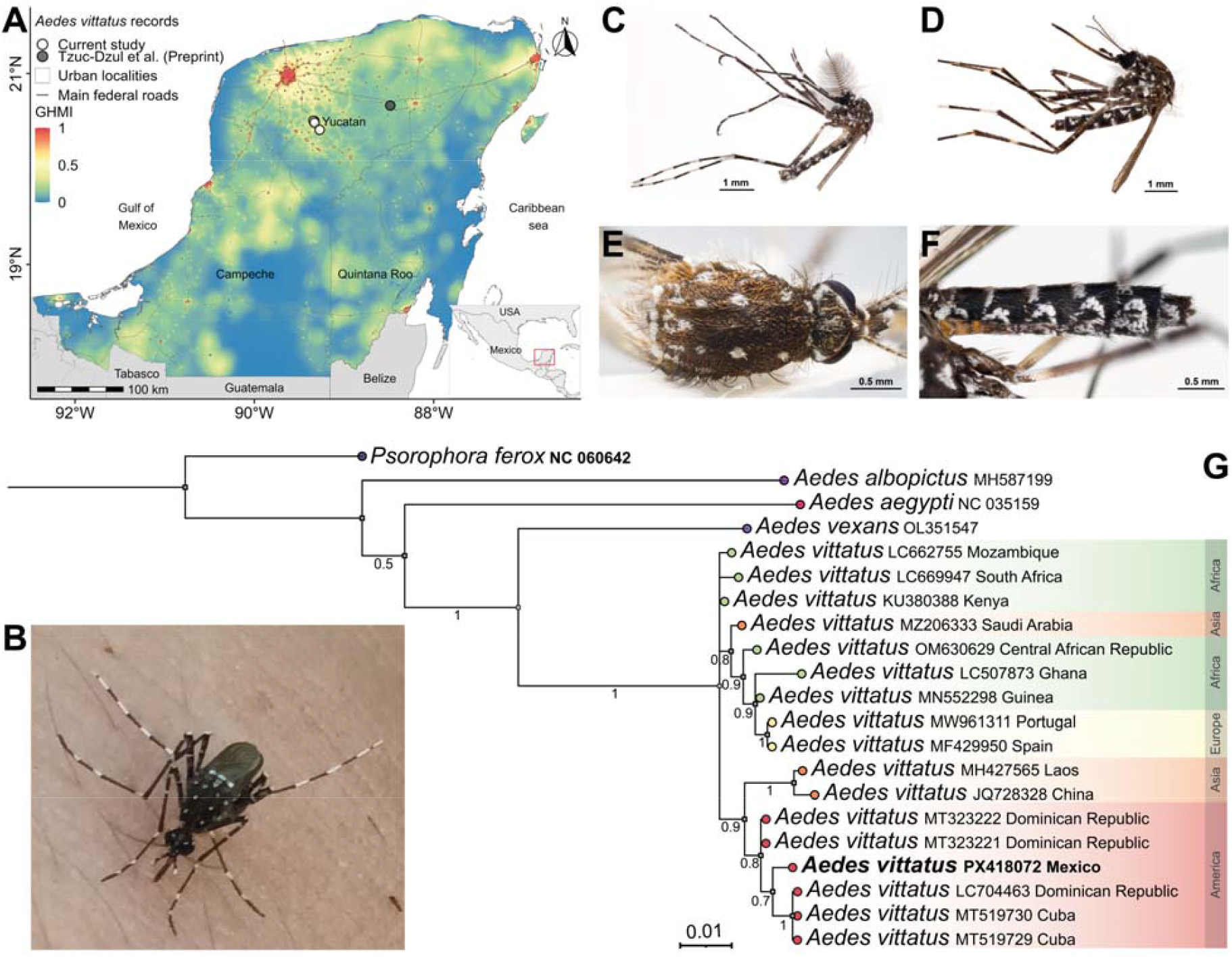
First confirmed population of *Aedes* (Fredwardsius) *vittatus* in continental America, Yucatán, Mexico. (A) Records of *Ae. vittatus* in the Yucatán Peninsula, showing the present study (white circles) and a previously reported specimen (gray circle; Tzuc-Dzul et al., preprint), overlaid on the Global Human Modification Index (GHMI; 0.09 km^2^ resolution). GHMI values range from 0 (unmodified) to 1 (completely modified). Gray lines indicate main federal and state roads. (B) Field observation of female *Ae. vittatus* landing on field staff in Mama, Yucatán, (C– F) Morphological characteristics of specimens of *Ae. vittatus*: (C) male lateral view, (D) female lateral view, (E) scutum with narrow white scale patches, (F) lateral view of abdomen. (G) Bayesian phylogenetic tree of COI sequences showing the placement of the Mexican specimen (bold, highlighted) within the American–Caribbean clade, clustering with sequences from the Dominican Republic and Cuba. Posterior probabilities are shown at nodes. Scale bars: C, D = 1 mm; E, F = 0.5 mm.

To confirm species identity, we sequenced a fragment of the mitochondrial COX1 gene from *Aedes* (Fredwardsius) *vittatus* collected in Yucatán, Mexico (genebank accession number PX418072), and analyzed it with global reference sequences. Bayesian phylogenetic inference placed the Mexican specimens within the American–Caribbean lineage, clustering with sequences from Cuba and the Dominican Republic (Figure 1G). Although the invasion history of *Ae. vittatus* is only beginning to unfold, this regional pattern resembles the early stages of *Aedes aegypti* expansion, for which the Caribbean acted as a bridgehead before dispersal into the Americas and beyond (8). While posterior support for the American *Ae. vittatus* subclade was moderate (0.8–0.9), the overall tree was well resolved, strengthening confidence in this inference. The case of *Ae. aegypti* illustrates how the Caribbean can serve as an intermediate launch point for Old World mosquitoes, underscoring the importance of acting now to monitor *Ae. vittatus* and prevent its wider establishment as a new invasive vector in the Americas.

We also characterized the ecological context using the Global Human Modification Index (GHMI; https://gdra-tnc.org/current/). High GHMI scores in the Yucatán Peninsula reflect intense land-use change from urbanization, agriculture, and infrastructure projects, highlighting conditions favorable for mosquito establishment and spread (Figure 1A). As a flat landmass with few natural biogeographic barriers, the peninsula provides little resistance to dispersal of habitat-tolerant invasive species. Studies of *Aedes aegypti* have shown that flat, highly connected regions with dense human activity enhance mosquito gene flow and facilitate spread (9). By analogy, regions where *Aedes* (Fredwardsius) *vittatus* is now reported—including the Yucatán Peninsula—present similar ecological and sociological conditions that may accelerate its population increase and dispersal, reinforcing the urgency of monitoring this emerging invasive species.

The detection of *Aedes* (Fredwardsius) *vittatus* in southeastern Mexico highlights the potential emergence of a new arbovirus vector in the Americas. The Yucatán Peninsula is undergoing profound anthropogenic change, where deforestation, agricultural expansion, and large-scale infrastructure projects such as the Tren Maya (10) are rapidly reshaping landscapes. Beyond their economic and social goals, such megaprojects can intensify ecosystem degradation, reduce ecological barriers, and enhance human connectivity, thereby creating ideal conditions for the establishment and spread of invasive mosquitoes. These dynamics underscore the need to integrate health considerations into land-use planning, recognizing that environmental transformation can amplify the risk of vector-borne diseases. Including *Ae. vittatus* in regional surveillance and control programs will be essential to anticipate its spread and mitigate future public health impacts.

## Supporting information

Appendix figure 1 and Methods for Molecular Identification of Aedes (Fredwardsius) vittatus

## Acknowledgments

We thank Fernando Chan-Poot for assistance with fieldwork and Humberto Bahena-Basave for photographing *Aedes vittatus*. We are also grateful to Marysol Trujano Ortega and Noemí Salas Suárez for their support with entomological laboratory materials at the ECOSUR Zoology Museum.

## About the Author

Dr. Chan-Chablé is a postdoctoral researcher at the Center for Research and Advanced Studies, Mérida Unit, Mexico. His research focuses on the natural history, taxonomy, and ecology of mosquitoes (Diptera: Culicidae) and other arthropods of public health importance in the Yucatán Peninsula.

## Notes

### Competing Interest Statement

The authors have declared no competing interest.

