## Appendix figure 1 and Methods for Molecular Identification of Aedes (Fredwardsius) vittatus for "Invasive *Aedes (Fredwardsius) vittatus* reaches Continental America"

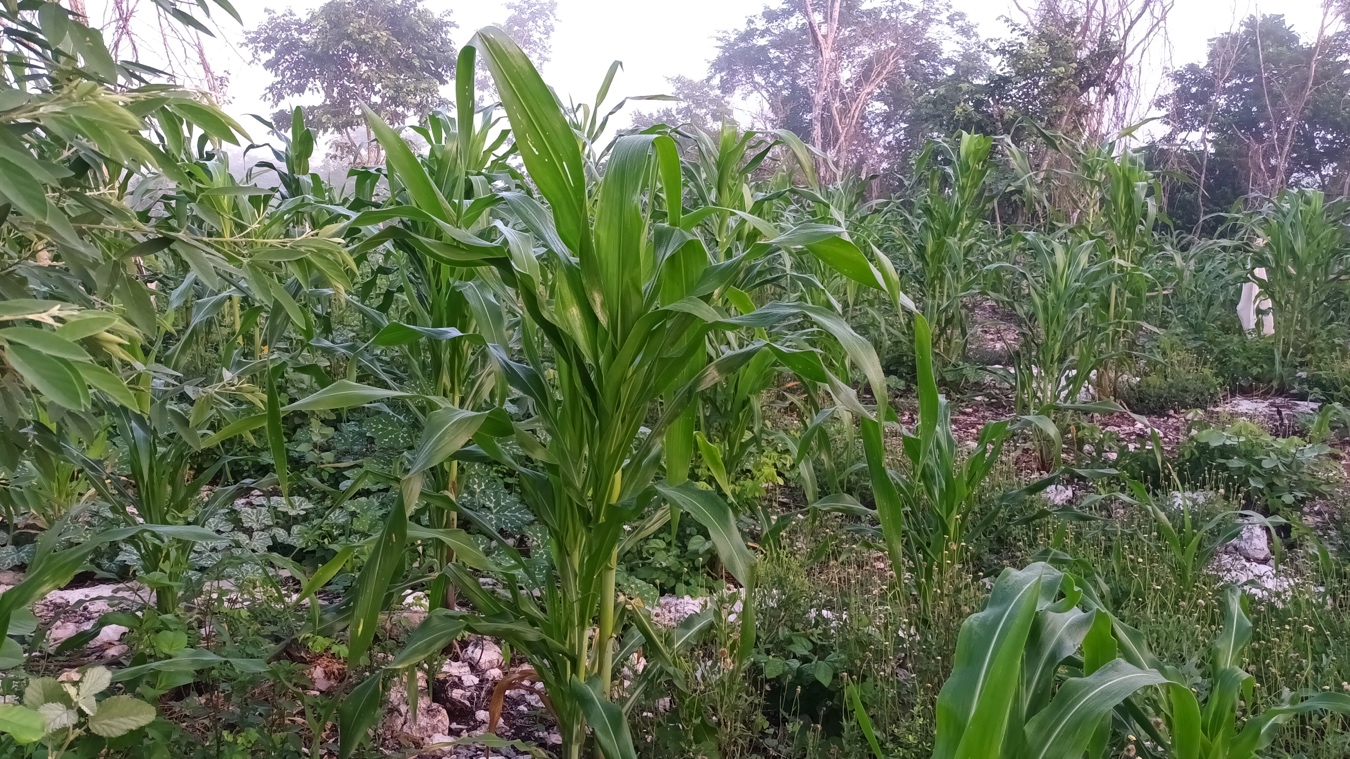

**Appendix Figure 1.** Ecological context of field observations of *Aedes* (Fredwardsius) *vittatus* in southeastern Mexico. Traditional Mayan cornfield (milpa) on the outskirts of Mama, Yucatán.

**Methods for Molecular Identification of *Aedes* (Fredwardsius) *vittatus***

**Nucleic acid extraction**

Genomic DNA was extracted using the Quick-DNA Miniprep (Zymo Research, USA), following the manufacturer’s instructions. Extractions were performed from mosquito legs. The final DNA elution was carried out in 50 µL of DNase/RNase-free water.

**Amplification of Cytochrome Oxidase 1 (COX1)**

PCR amplifications were performed in a final reaction volume of 25 μL, consisting of 12.5 μL of Phusion High-Fidelity DNA Polymerase (Thermo Fisher Scientific, USA), 1 μL of each primer (10 μM), 7.5 μL of ultrapure water, and 3 μL of genomic DNA. All reactions were conducted in a Veriti™ 96-well thermal cycler (Applied Biosystems™, Foster City, CA, USA).

A fragment of the COX1 gene was amplified using primers LCO-1490 (5′- GGT CAA CAA ATC ATA AAG ATA TTG G - 3′) and HCO-2198 (5′- TAA ACT TCA GGG TGA CCA AAA AAT CA - 3′) (1). PCR consisted of an initial denaturation at 94 °C for 1 min; followed by 39 cycles of denaturation at 94 °C for 15 s, annealing at 48 °C for 30 s, and extension at 72 °C for 45 s; with a final extension at 72 °C for 7 min.

Amplicons from PCR assays were resolved by electrophoresis on 1% agarose gels prepared in 1× TAE buffer, stained with RedGel (Biotium, USA), and visualized under UV light using a BioDoc-It2™ imaging system (Analytik Jena US LLC, USA). PCR products were sequenced by Sanger sequencing.

**Bioinformatic analyses**

Forward and reverse reads were assembled and edited to generate consensus sequences using Geneious Prime v 2025.2.2 (2). Sequence alignments were performed using the MAFFT online server v7 (https://mafft.cbrc.jp/alignment/server/) under default parameters (3). The resulting alignments were manually reviewed and edited in AliView v1.28 (4). The best-fit nucleotide substitution model was selected using jModelTest2, implemented through the CIPRES Science Gateway (https://www.phylo.org/) with default settings (5).

Mitochondrial COX1 marker sequences of Aedes spp. were retrieved from GenBank (accession numbers shown in Figure 2) and include representatives from Africa, Asia, Europe, the Caribbean, and the newly generated Mexico sequence.

Phylogenetic relationships among *Aedes vittatus* sequences were inferred using Bayesian Inference (BI) in Geneious Prime v 2025.2.2, applying the GTR + G substitution model and designating *Sorophora vorax* as the outgroup (6). Markov Chain Monte Carlo (MCMC) analyses were run for two million generations, with trees sampled every 100 generations. Posterior probabilities were estimated from the sampled trees after discarding the first 25% as burn-in to ensure convergence and stationarity. The final phylogenetic tree was visualized and edited using FigTree v1.4.4 (7).
